# Structural Learning of Proteins Using Graph Convolutional Neural Networks

**DOI:** 10.1101/610444

**Authors:** Rafael Zamora-Resendiz, Silvia Crivelli

## Abstract

The exponential growth of protein structure databases has motivated the development of efficient deep learning methods that perform structural analysis tasks at large scale, ranging from the classification of experimentally determined proteins to the quality assessment and ranking of computationally generated protein models in the context of protein structure prediction. Yet, the literature discussing these methods does not usually interpret what the models learned from the training or identify specific data attributes that contribute to the classification or regression task. While 3D and 2D CNNs have been widely used to deal with structural data, they have several limitations when applied to structural proteomics data. We pose that graph-based convolutional neural networks (GCNNs) are an efficient alternative while producing results that are interpretable. In this work, we demonstrate the applicability of GCNNs to protein structure classification problems. We define a novel spatial graph convolution network architecture which employs graph reduction methods to reduce the total number of trainable parameters and promote abstraction in intermediate representations. We show that GCNNs are able to learn effectively from simplistic graph representations of protein structures while providing the ability to interpret what the network learns during the training and how it applies it to perform its task. GCNNs perform comparably to their 2D CNN counterparts in predictive performance and they are outperformed by them in training speeds. The graph-based data representation allows GCNNs to be a more efficient option over 3D CNNs when working with large-scale datasets as preprocessing costs and data storage requirements are negligible in comparison.

## 1. Introduction

Traditional machine learning approaches have been extensively applied to numerous problems in structural biology. Methods such as support vector machines (SVMs), artificial neural networks (ANNs), and Random Forest (RF) have been used for protein classification [1], fold recognition [2, 3, 4, 5, 6, 7, 8], secondary and tertiary structure prediction [9, 10, 11, 12, 13], protein-protein interaction prediction [14, 15], protein-ligand binding affinity prediction [16, 17, 18, 19, 20, 21, 22], scoring of computationally generated protein models (decoys) [23, 24, 25, 26] and decoys quality assessment [7, 26, 27, 28, 29, 30, 31]. A major drawback of these methods and traditional machine learning methods as a whole is their dependence on hand-engineered features. Feature selection greatly impacts model performance [32], but defining appropriate feature sets for any given task requires a substantial amount of domain-expert knowledge and heuristics.

Recently, deep learning methods and in particular convolutional neural networks (CNNs) methods have popularized for their ability to perform seamless feature extraction on structural data without the need of domain knowledge. 3D CNNs have become a standard approach to deal with structural data through architectures like VoxNet [33] and ShapeNet [34], and consequently their application to structural biology has become a very active area of research. To be fed into these networks, the 3D structures of molecules are represented on a grid and a classification or regression problem is solved by using only these grids and in some cases a few additional channels of information. For example, Derevyanko et al. developed a 3D CNN-based approach to score and perform quality assessment (QA) of decoys using their 3D atomic densities only [35, 36]. Li et al. proposed a 3D CNN-based approach to perform quality assessment of RNA 3D structures using a 3D grid representation of the structure as input and without extracting features manually [37]. 3D CNNs have been applied to the prediction of protein binding sites and their interactions with ligands [38, 39, 40, 41]. Also, they have been used for structure classification tasks [42].

The aforementioned methods use 3D volumetric representations of molecular structures to learn features of a molecule through its shape, more specifically how the structure occupies 3D space. However, there are several limitations when applying these approaches to structural biology. Firstly, there is a large computational overhead associated with generating and storing coarse-grained images of molecular structures, especially at resolutions which produce biologically meaningful features. In addition, image resolution is highly dependent on the size and shape of structures requiring a number of preprocessing steps to ensure all structures are effectively captured within the image window. Secondly, volumetric images of molecular structures are sparse while convolution-based neural networks are best fit for dense inputs. As a result, training on sparse data using 3D CNNs is severely inefficient, an effect of ‘empty’ convolutions being applied over large spans of unoccupied space in high resolution images. Architectures such as OctNet [43] have been developed to deal with sparsity of volumetric images, though the method requires additional preprocessing to convert volumetric images into the algorithm’s tree-based data structures. Furthermore, 3D CNNs are not rotationally invariant meaning the method requires either substantial rotation-based data augmentation or additional preprocessing in the form of orientation normalization. Finally, the computational cost is compounded when chemical and other biologically significant properties are incorporated as additional channeled information within the representations.

2D CNNs offer an alternative, cost-effective approach through representations or transformations that capture some aspects of the 3D structure in the 2D space. Corcoran et al. [44] apply space filling curves to convert 3D structures to 2D and then use a 2D CNN for protein classification achieving results comparable to those achieved by the 3D CNN counterpart with decreased training time cost. Tavanaei et al [45] classify proteins encoded by tumor suppression genes (TSGs) and protooncogenes (OGs) from 2D feature map sets extracted from their 3D structures. Wang et al. [46] achieve de novo prediction of contact maps by using ultra deep 1D and 2D CNNs. Zacharaki [47] outperforms her 3D CNN-based results [42] for the classification of an enzyme dataset by encoding amino acid interactions through the distribution of pairwise amino acid distances. However, although these methods are effective at capturing structural fingerprints that seem to be enough for protein classification tasks, it is impossible to connect those fingerprints to biologically relevant features, which is key to feature extraction. In fact, learning how the networks work and interpreting the results they produce may elucidate meaningful characteristics that contribute to the classification task and that were not evident to the domain scientists.

A less explored approach is offered by graph-based convolutional networks (GCNNs), which allow for a natural spatial representation of molecular structures without the computational cost of volumetric representation or the interpretability issues of the 2D representations. To the best of our knowledge, they have not yet been widely applied to the field of structural biology where they can learn local and global features by integrating information across graphs that represent the three dimensional structure of molecules. For example, Fout et al.[48] proposed a graph convolution approach for the prediction of protein-protein interactions. The success of graph convolution in protein-protein interaction prediction allows for the possible expansion of similar graph convolution operations to other protein structure classification and regression tasks, such as protein function prediction, protein decoy scoring, and protein-ligand binding affinity prediction, although a clear definite formulation of spatial graph convolutions for micro and macro molecular problems is still an active area of research.

In this paper, we discuss the applicability of GCNNs to protein classification. We define a novel spatial graph convolution network architecture that employs graph reduction methods to reduce the number of training parameters and promote abstraction in intermediate representations. We explore the efficiency and performance of GCNNs and compare them to 2D CNNs on a wide range of protein structure classification problems including structural classification of cancer-related proteins, functional classification of enzymes, active/inactive conformation classification of kinases, and classification of a subset of the RAS protein family into KRAS and HRAS isoforms.

We show that GCNNs are able to learn effectively from simplified graph representations of protein structures thus offering a more interpretable approach for structural data when compared to its 3D and 2D CNN counterparts. Simplified data representation allows GCNNs to be a more efficient option over 3D CNNs when working with large-scale datasets as preprocessing costs and data storage requirements are negligible in comparison. They can seamlessly operate on proteins of any size and require less data augmentation as standard 3D CNNs thanks to the GCNN’s rotational invariance. Our results are comparable to those achieved by the 2D CNN approach in [47] in terms of predictive performance but they are worse than the 2D CNN method in training speeds. However, they show an advantage when it comes to interpretability.

## 2. Methods

### 2.1 Dataset

In this work, we use four different protein datasets from recent studies for protein structure classification problems. All protein structures used for this work were retrieved from the RCSB Protein Data Bank (https://www.rcsb.org) [49] in PDB formatted files.

### 2.1.1 Datasets for Protein Classification

The four selected datasets evaluate our method’s ability to detect structural differences at various granularities (i.e globally different, globally similar/locally different), to process sequentially and structurally different structures, to assess the scalability of our method, and to process protein structures with multiple chains. A short description of each dataset is listed below, and the details of each dataset’s class composition are summarized in Table 1.

**Table 1:** Number of examples in each class for K-Ras/H-Ras, TSG/OG, Active/Inactive Kinase, and Enzyme datasets.

|  | Class 1 | Class 2 | Class 3 | Class 4 | Class 5 | Class 6 | Total |
| --- | --- | --- | --- | --- | --- | --- | --- |
| KRas/HRas | 77 | 156 | - | - | - | - | 233 |
| TSG/OG | 1,176 | 1,138 | - | - | - | - | 2,314 |
| Kinase | 1,773 | 1,592 | - | - | - | - | 3,365 |
| Enzymes | 11,194 | 18,733 | 23,861 | 4,476 | 2,762 | 2,532 | 63,558 |

- **K-Ras/H-Ras** Provided by Corcoran et al. [44], this dataset is composed of 233 structures belonging to two sub-classifications of the Ras protein, K-Ras and H-Ras. The Ras family of proteins are of great interest in cancer research [50, 51] as mutations of proteins in this family are thought to contribute to the development of cancers such as pancreatic cancer and colorectal cancer. Structures in this dataset are on average 170 amino-acids in length and 59.92Å wide in diameter.
- **TSG/OG** - Provided by Tavanaei et al. [45], this dataset is composed of 2,314 structures belonging to two different classes of proteins, tumor suppression genes (TSGs) and proto oncogenes (OGs), whose regulatory mechanisms play major roles in the suppression and growth of tumors respectively. Identification of structures with similar functionality to TSGs and OGs is valuable to the development of cancer treatments, which target these two proteins. Structures in this dataset are on average 289 amino-acids in length and 64.48Å wide in diameter.
- **Active/Inactive Kinase** - Provided by McSkimming et al. [52], this dataset consists of 3,365 kinase protein structures with either active or inactive conformations. Determining the activity of kinase structures is valuable in understanding the regulatory mechanism of kinases. While very similar in overall structure, subtle differences between the structures’ binding site conformation is the main determinant of kinase activity. Structures found in this dataset are on average 300 amino-acids in length and 58.62Å wide in diameter.
- **Enzymes** - Provided by Amidi et al. [42], this dataset is the largest of the four containing a total of 63,558 structures belonging to 6 different enzyme classifications according to the Enzyme Commission (EC) scheme. More specifically, these structures belong to EC classifications 1 through 6. Unlike the previous datasets, structures within this set are composed of one or more protein chains. They are on average 968 amino-acids in length (total number along all chains) and 75.04Å wide in diameter.

## 2.2 Data Representation

Data representation plays an important factor when deciding a suitable deep learning approach. Different network architectures take advantage of inherent structures present within their input data to efficiently reduce dimensionality and promote forms of invariance. For example, traditional CNN architectures are able to learn translationally invariant feature representations by exploiting the uniform lattice structure of 2D images, learning locally connected feature detectors called ‘kernels’, and applying them along both axes of the image. In this work, we compare our graph representation method for protein structures to a previously used data representation method based on statistical profiles.

### 2.2.1 2D Statistical Profiles

Work done by Zacharaki et al. [47], has shown that structural classification of proteins can be performed using statistical profiles of the density of pairwise amino-acid distances within a structure. 2D images from these pairwise statistics were generated and used to train 2D CNNs with success, although a good intuition as to why their particular architecture learned effectively from this representation is missing from their exploration. They constructed these statistical profiles by calculating the distances between each amino acid type for proteins in their dataset. These distances were found using the coordinates of the alpha-carbon in each amino-acid. For each amino-acid pairwise combination, the distances were standardized over a histogram of equally-sized bins. A total of 9 bins were used for each histogram, with distance values ranging from 0 to 50Å. Thus, the generated feature maps are of shape *NxNx*9, where *N* is the number of amino-acid types. Using the 23 different kinds of amino acids defined in the PDB format, the final shape of a protein’s 2D feature map is 23*x*23*x*9.

## 2.3 Graph Representation of Proteins

In machine learning, we are often confronted with structured data, which can be easily represented as graphs such as social networks, knowledge graphs, and molecules [53]. In the case of protein structures, an intuitive graph representation of a protein can be formed by considering amino-acid monomers or residues as distinct nodes whose edge connections describe spatial relationships between them. In general, a graph *G* is defined by tuple (*V, A*), where *V* ∈ ℝ^*NxF*^ defines the vertices or nodes of the graph, *N* the number of nodes and *F* the number of features in each node. The adjacency matrix *A* ∈ ℝ^*NxN*^ defines edge connections between *N* nodes, where *A*_*ij*_ is a scalar weight value representing the strength of relation between nodes *i* and *j*. Using this definition of a graph, protein structures can be defined as graphs whose residue features are encoded in node features *V* and spatial proximity between residues are encoded in adjacency matrix *A*.

### 2.3.1 Constructing Node Features

To construct node features *V* for a given protein, we first gather 4 features for each node as defined in the PDB file: 1) the type of residue that the node represents, 2) the residue’s positional index along the primary structure, 3) the residue’s *Cα* (*x, y, z*) coordinates, and 4) the residue’s side-chain (*x, y, z*) center-of-mass coordinate. Using 3 and 4, two other structural features are derived: 1) the residue depth with respect to the structure’s center-of-mass and 2) the orientation of the side chain with respect to the structure’s center of mass. The process used to encode residue type, position, depth and side-chain orientation to the final node feature tensor *V* is described below.

First, the residue types were encoded as one-hot vectors forming residue features *V*_*res*_ ∈ ℝ^*Nx*23^. One-hot vectors are a common way of representing categorical values in machine-learning where the occurrence of a specific category is denoted as 1 with the rest of categories denoted as 0. All 23 residue types defined in the PDB format were considered for this approach resulting in the *Nx*23 dimensionality of *V*_*res*_.

Second, the index of each node was encoded using the sinusoidal positional encoding described in [54]. This approach is commonly used to encode position of words in sentences for use in transformer-based networks, and was used here to encode the primary structure’s sequential information as an added node feature. This allows features to be learned from the relative position of a node along the proteins’ primary structure. The positional encoding is defined through functions

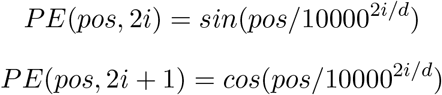

where *pos* is the positional index and *i* is the encoding dimension. The number of *d* dimensions used for the positional encoding depends on the length of the largest protein, though we found that 4 encoding dimensions were sufficient for all classification problems. The positional encoding of the residues form positional feature tensor *V*_*pe*_ ∈ ℝ^*Nx*4^.

Third, the relative depth of the residue with respect to the center of mass was calculated for each residue. This was done by normalizing the distance between the residues’ *Cα* atom and the center of mass of the structure over all nodes in the structure, giving the feature values between [0.0, 1.0]. This feature provides information about the packing properties of the structure, which is a key characteristic needed to understand differences between proteins [55]. The residue depth features form feature tensor *V*_*depth*_ ∈ ℝ^*Nx*1^.

Fourth, the orientation of the side chain with respect to the center of mass was calculated by finding the cosine similarity between two vectors originating from the residues *Cα* atom: 1) the vector to the center of mass and 2) the vector to the center of mass of the side chain. This feature encodes the orientation of the side-chain to values between [−1.0, 1.0] and represents information about the hydrophobic properties of each residue in the structure. Side-chains oriented towards the center of the protein are typically hydrophobic (water-hating) while side-chains oriented away from the center of the protein are typically hydrophilic (water-loving). The side-chain orientation form feature tensor *V*_*orien*_ ∈ ℝ^*Nx*1^.

Finally, *V*_*res*_, *V*_*pe*_, *V*_*depth*_, and *V*_*orien*_ were concatenated along their feature axis resulting in the final node feature tensor *V* ∈ ℝ^*Nx*29^ for each protein graph. The 3D coordinates for each residues’ *Cα* atom were then gathered into coordinate tensor *C* ∈ ℝ^*Nx*3^. All node coordinates for a given structure were normalized by moving the center-of-mass to the origin (0, 0, 0).

Due to the varying length of proteins’ primary structure within a dataset, node features and coordinates must be formatted to fit the majority of the training examples. For this reason, *V* and *C* were both either zero-padded or truncated accordingly in order to fit the average length protein within a dataset. In terms of preprocessing, *V* and *C* are the only required inputs to our model, but further processing is conducted internally within our network architecture to generate the required edge matrices prior to performing graph convolution.

Often times, residues are missing from a protein’s PDB data due to issues when measuring the experimentally determined structure of that protein. In order to deal with this problem of missing data in our graph representation, we include the missing residues with only amino-acid and positional encodings, and use the *Cα* coordinates of the last existent residue along the sequence to reduce noise when performing graph reduction. During training, a mask is also provided to indicate not to convolve over missing or empty residues in order to reduce the noise during convolution.

### 2.3.2 Constructing Edge Features

As expressed in the previous section, the construction of the adjacency matrix tensor *A* ∈ ℝ^*NxN*^ is conducted within the network framework prior to convolution. This allows for the construction of the graphs to be accelerated using GPU hardware alongside the model. While increasing the total computational cost of the model, the amount of resources needed to store and load protein structural data becomes negligible as a result. Within the framework, the Euclidean pairwise distance between each node in coordinate tensor *C* is calculated using

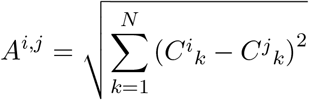

In order to perform graph convolution, these pairwise distance must be regularized in order to promote stable gradient descent. Our method of normalizing these pairwise distances within our network architecture is further discussed in section 2.6.2.

## 2.4 Data Augmentation

Although no augmentation strategy was used for the 2D statistical profile data in work done by Zacharki et. al, a method was developed for graph data used in this study. The following section describes that data augmentation strategy.

### 2.4.1 Graph Datasets: Random Perturbation of Node Positions

Unlike 2D and 3D CNNs, graph convolutions are rotationally invariant and consequently, there is no need to supplement the training data with rotation-based augmentations. Nevertheless, as with any deep learning approach, more training data allow models to better converge on a solution. One augmentation strategy explored for augmenting graphs was to generate perturbations of the structure where each node is shifted in position according to a random normal distribution along a set radius. The number of augmentations and the radius used for the random perturbation were defined according to the augmentation needs of each dataset.

While this method of data augmentation shows to help improve gradient descent and prevent over-fitting on small datasets, this type of random shifting does not take into account transformations that may result in biologically impossible structures. For example, two residues may be shifted into close proximity to one another, which, in the real world, may not be possible due to repulsive forces between the two bodies preventing such configuration. As a result, it may be more appropriate to generate augmentations based on molecular dynamics simulations to prevent such configurations and provide more biologically meaningful augmentations, though this in itself would require a large amount of computational resources and cannot be done in-line with training. That being said, augmenting data by providing valid variations of structure would be impractical when training 3D CNNs as you would most likely need to supplement the perturbations with additional rotational augmentations.

## 2.5 Deep Learning Approaches

One of the most popular deep learning methods is the convolutional neural network (CNN), which utilizes shared local filters and hierarchical information processing analogous to the brain’s visual system. While CNNs deal only with inputs defined as regular 2D or 3D grids, GCNN architectures apply to graphs that represent irregularly structured data. Different graph convolution implementations have been proposed [56] each tackling a specific subset of possible graph structures. We derive our implementation from previous work conducted on a spatial variant of graph convolutions [56]. In the following sections, we briefly describe traditional CNNs, and then describe our implemented spatial GCNN architecture in more detail.

### 2.5.1 CNNs

CNN-based systems have made substantial breakthroughs in feature extraction and image recognition tasks [57]. CNNs are able to learn hierarchies of features by convolving a large number of randomly-initialized feature detectors, known as ‘kernels’ over their input data. These kernels look at small subsections of an image, and are sensitive to various combinations of pixel clusters, activating in the presence of learned features. Each layer of a CNN learns an increasingly abstracted representation of the input data through a combination of convolution and feature reduction (pooling) operations, with earlier layers learning low-level features such as edges or curves and later layers learning high-level features such as shapes or textures. The final layers of the network contain a highly-distilled set of features which can in turn be used to form classifiers and regressors. The learning capability of CNNs, as in all other neural network systems, is enabled through backpropagation algorithms, which provide for the efficient propagation of error gradients throughout the layers of a network in response to the network’s performance defined in its loss function. This error gradient is then used to adjust the weights of each layer of the network in order to “zero-in” on a high-performing configuration at the parameter level.

One way of intuitively understanding the benefits of this style of architecture over classic artificial neural networks (ANNs) is through the idea of parameter reuse. Since we can assume that there exists some regularity between local features in certain kinds of images, this regularity can be exploited in order to reduce the total amount of trainable parameters needed in a network. Within CNN architectures, kernels in themselves are small locally-connected neural networks whose weight parameters can be applied along any portion of the input image space. Thus, trained CNN-style kernels can be used to detect the presence of a feature in any part of an image even if that feature has been shifted along an axis. This ability to exploit inherent regular structures in data helps reduce the parameter complexity of models, allows for much deeper architectures due to the reduced computational overhead, and overall improves generalizability through model invariance.

3D CNNs are a straightforward extension of 2D CNNs and differ only in that they convolve over an added depth dimension. This style of networks has become a popular method of learning from volumetric images and videos [58], though they are substantially more expensive to run than 2D CNNs. The added dimension means the time complexity of 3D CNNs scales cubically with the size of the input images. This results in a computational resource bottleneck when working with high-resolution images, images with a large number of channels, and deeper network architectures. Methods have been explored to reduce the cost of performing convolution on 3D images such as methods that reduce the number of ‘empty’ convolutions [43] and methods for 3D image dimensionality reduction that allow optimized 2D convolution implementations to be used instead [59, 44]. We decided on excluding 3D CNNs from this study due to the high computational overhead of this approach.

### 2.5.2 Spatial Graph CNN

Spatial graph convolutions are a subdomain of graph convolutions, which mainly work with non-spectral methods of parameterization [56]. Non-spectral approaches define convolutions directly on the graph, operating on ‘spatially’ close neighbors (k-hop nearest neighbors). The major challenge of non-spectral approaches is defining a convolution that is able to operate on differently sized neighborhoods while maintaining local invariance of the network. As for the pairwise graphs used to define protein and molecular structures in this study, differently sized neighborhoods were not an issue, though the dense pairwise representation gave rise to three new problems: 1) edges represent distances in Euclidean space not local connectivity, 2) local neighborhoods (1-hop connections) contain all nodes, and 3) the independent parameterization of each edge results in an extremely sparse and inefficient operation using current implementations of matrix multiplication in neural network libraries such as Tensorflow [60]. The following subsections detail the main operations used in our GCNN architecture to tackle these issues as well as how we perform graph reduction in order to facilitate the construction of deeper architectures and the processing of larger graphs. Figure 1 shows a diagram of the full GCNN architecture.

**Figure 1:**
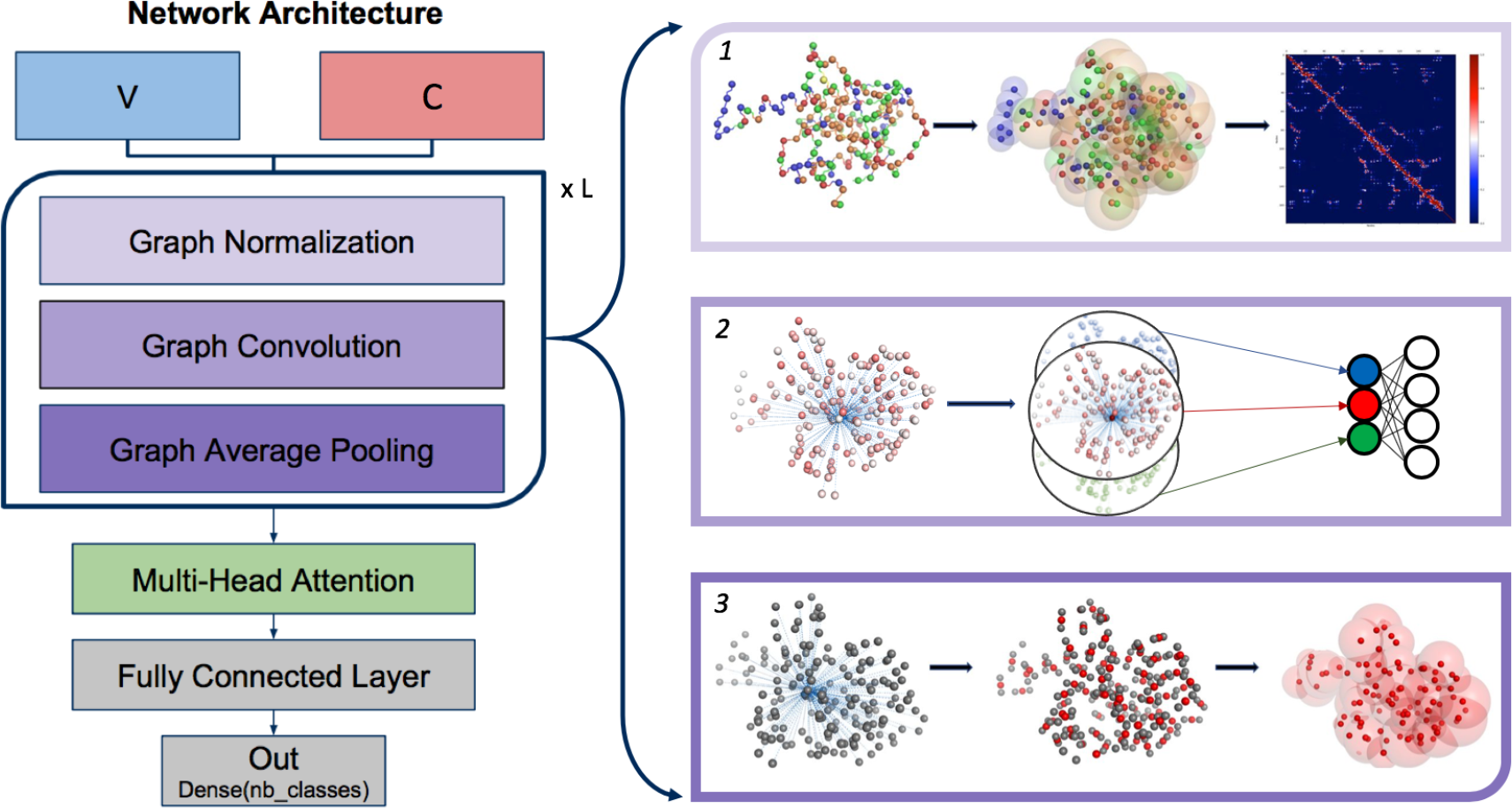
A diagram of the spatial GCNN architecture. The model takes in input tensors V and C which are operated on sequentially by *L* number of graph convolutional blocks. This is followed by a layer of multi-head attention, a fully connected layer of neurons, and finally the output classification prediction. The right of the diagram provides visualizations of the three operations forming a graph convolutional block in top-down order.

## 2.6 Spatial Graph CNN Implementation

### 2.6.1 Performing Graph Convolution

Our graph convolution operation is based of Such et al.’s spatial graph convolution [56]. To start, we take a tensor of pairwise adjacencies *A*′ ∈ ℝ^*NxMxN*^ and a tensor of node features *V* ∈ ℝ^*NxF*^ and output a new tensor of convolved node features *V*′ ∈ ℝ^*NxK*^, where *M* is the number of total adjacency matrices, *N* is the number of nodes in the graph, *F* is the number of features in each node, and *K* is the number of new features after convolution. Convolving over a node involves a two-step process. First, the features of neighboring nodes are propagated to the convolved node, and then the accumulated features are mapped to a new set of features using a fully connected neural network. Convolution over a graph is more specifically defined as

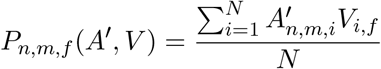

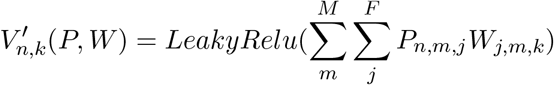

where *P* ∈ ℝ^*NxMxF*^ is the propagated node features along all adjacency matrices, and *W* ∈ ℝ^*FxMxK*^ is a trainable weight tensor. While straightforward and easily implementable using a few dense matrix multiplications, the issue with this method is evident when it comes time to fully parameterize or assign weights to all graph edges independently from one another. Doing so would require each edge column to be placed in independent adjacency matrices resulting in a tensor *A*′ of dimensionality *NxNxN*, making it very expensive to independently parameterize all pairwise relationships within a graph.

Without another form of parameterization or method of performing efficient sparse matrix multiplications, we are limited to assigning a single trainable parameter to all neighboring node connections over a given convolved node or arbitrarily dividing edges into separate adjacency matrix according to some heuristic. The next section describes our method of graph normalization, which in addition also acts as a method of improving edge parameterization while remaining computational feasible overall.

### 2.6.2 Normalizing Graph Using Learned Gaussian Kernel

To learn effectively using graph convolutions, edges of the constructed graphs must be normalized to allow for proper feature communication between nodes. More specifically, edge values should lie between [0.0, 1.0] where 0.0 indicates that no information will be passed between nodes and vice-versa. The normalization strategy should also capture some aspect of spatial locality, where nodes closer together in Euclidean space would be considered to be more related. To exhibit this behavior we used the Gaussian function

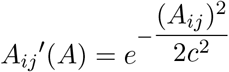

where *A* is our pairwise distance matrix and *c* is the Gaussian RMS width. Parameter *c* allows us to control the radius of the Gaussian bell curve which in turn can be used to define the limits of a local neighborhood around a given node in Euclidean space. Initial exploration found that a good starting value for *c* is the average diameter D of all the structures in the dataset, though this is far from an optimal value. Furthermore, this method assigns a global static value for *c* and does not differentiate between nodes with different sets of feature vectors.

Given a node’s feature set, different values of *c* may more appropriately define the optimal limit for its local neighborhood. To learn the best neighborhood radius for a node given its feature set, we formulated a training procedure to learn parameter *c*

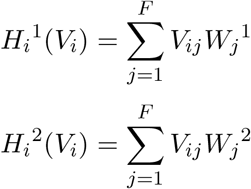

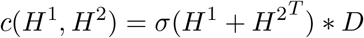

where *W*^1^ ∈ ℝ^*F*^ and *W*^2^ ∈ ℝ^*F*^ are trainable weight tensors and *σ* is a sigmoid activation. The two neural networks are applied along all nodes, and are then combined through transpose addition of the two output tensors. This allows for a unique value to be derived for each possible pairwise combination in the constructed graph. Sigmoid activation is then applied and then multiplied with the average structure diameter to scale it between the values of (0.0, *D*]. The resulting values are then applied in line with the Gaussian kernel on a per edge basis.

Our implementation of this normalization strategy allows for any number of Gaussian filters to be learned in parallel. These Gaussian filters can be used to construct a set of normalized adjacency matrices which can then be subjected to graph convolution. This method provides a way of parameterizing the graph convolution by learning multiple permutations of possible local neighborhoods, which can be convolved over. Further investigation must be conducted to prove that this method can be used to approximate full parameterization as discussed in the previous section.

### 2.6.3 Reducing Graph Size Using Sequence-Based Averaging

Performing graph convolution on large graphs becomes very expensive when running deeper networks. In order to be able to process large graphs while still maintaining deep network architectures, we define a method of reducing protein graphs by taking advantage of the inherent sequential ordering of the data. When constructing the graphs, nodes in feature and coordinate tensors *V* and *C* are ordered according to the primary sequence of the protein. It can be assumed that neighboring nodes are also close in spatial proximity. Based on this assumption, a quick and computationally efficient graph reduction strategy can be employed by averaging node feature values and coordinates along the sequence of the protein.

To perform a graph pooling, 1D average pooling is first applied to both tensors *V* and *C*. After this, the pooled *C* is used to construct a new set of pairwise adjacencies for the reduced graph. This method of graph reduction is able to preserve the overall global backbone structure of the protein. During development, max pooling was also explored for the reduction of feature tensor *V*, but resulted in less stable gradient descent when compared with average feature pooling.

### 2.6.4 Promoting Shift Invariance Using Multi-Head Attention

While developing our interpretability methods, we found a fail-case of an earlier GCNN architecture when alignment permutations of the proteins were fed for inference. Examples were truncated and shifted as to align all the structures along a common set of starting residues, and this resulted in model predictions with losses much higher than those seen during testing. This issue is thought to be due to the fully connected layer being sensitive to shifts in the ordering of nodes of the graphs. In addition, the graph convolutional block would produce different intermediate representations than those seen during training as a result of noise introduced by the graph reduction method applied on the shifted structures.

**Table 2:** Number of trainable parameters for each network on the 4 classification tasks. Total count include bias parameters. Smallest model per dataset are highlighted in boldface.

| Model | Dataset | Trainable Parameters |
| --- | --- | --- |
| 2DCNN | KRas/HRas | 505090 |
| GCNN | KRas/HRas | **124194** |
| 2DCNN | TSG/OG | 505090 |
| GCNN | TSG/OG | **266814** |
| 2DCNN | Kinase | 505090 |
| GCNN | Kinase | **217138** |
| 2DCNN | Enzyme | 505090 |
| GCNN | Enzyme | **378370** |

In order to provide a regularized ordering of intermediate features, we employed a layer of multi-head attention which is a common component of current state-of-art natural language translation models [54]. Attention works by detecting the presence of contextually relevant features along a sequence elements and combine those features into a new sequence of elements according to a learned weighted contribution. Previous work has been conducted using similar attention mechanisms to sort sequential data [61].

More investigation is required to understand the full drawbacks and benefits of using this operation in our model, though current results show that the use of multi-head attention resolves the issues found during our interpretation analysis. We believe this operation may also be helpful in regularizing the detection of secondary structure formation between examples, whose structure often vary in exact starting and ending residues.

## 2.7 Network Architectures

This section describes the network architectures explored for both types of data representation methods as well as the loss function and optimizer used to form the classification tasks. A summary of each network’s number of trainable parameters (including biases) can be found in Table 2.

### 2.7.1 2D CNNs

Network design for our 2D CNN models was derived from the work conducted by Zacharaki et al. [47]. Their particular architecture was relatively simple by current standards in image recognition using CNNs with no use of any non-sequential operations such as those found in residual CNNs (ResNets)[62] or densely-connected CNNs (DenseNets)[63]. The final network architecture for each individual CNN contains 2 convolutional layers operating with kernels of size (3×3), strides of (1×1), 128 filters and use Relu as the activation function. Each convolutional layer is then followed by a max pooling layer of kernel size (2×2) with dropout. These layers are followed by a single fully-connected layer with 512 neurons with dropout applied to both. Batch normalization was also employed for both convolution layers and fully-connected layer, and was applied prior to the activation functions. This is then followed by the final output layer with the number of neurons equal to the number of different classes. Softmax activation is applied to final outputs to get the predicted class probabilities. This network architecture was used for all 4 datasets as no changes needed to be made to accommodate the input data.

### 2.7.2 GCNNs

Network architectures for GCNN models vary according to the requirements of each dataset. Our GCNN architectures employ a number of convolutional blocks, which apply graph normalization, graph convolution, and graph pooling in that order. The number of convolutional blocks was found to be dependent on the overall size of the input graphs, with larger graphs requiring more blocks for dimensionality reduction. Through hyper parameter exploration, we found that network performance diminishes with graph pooling factors larger than 4 on the smaller protein datasets while still preforming considerably well on the Enzyme dataset up to a factor of 10. The number of learned Gaussian kernels and graph convolution filters were found to depend on the classification tasks difficulty, increasing according to the difficulty of the problem. After the graph convolution blocks, each architecture variant is followed by a layer of multi-head attention with heads equal to the number of nodes from the last convolutional block and a fully-connected layer of 128 neurons with dropout. This layer, like in the previous networks, is then followed by the final output layer with softmax. We found that the architecture works well using Leak Relu as the activation function for the convolutional and fully connected layers. In addition, batch normalization was employed prior to activation in those same layers.

### 2.7.3 Loss Functions and Optimizers

We chose categorical cross entropy [57] as the loss function for network training as it is commonly used for both binary and multi-class classification tasks. The ADAM optimizer [64] was chosen to minimize training loss as it provides computationally efficient back-propagation and tends to converge on solutions more quickly than standard stochastic gradient descent (SGD).

## 3. Results

### 3.1 Predictive Performance for the Protein Classification Study

For our training experiment, we split each dataset into training, validation and test sets according to a 70%/10%/20% random split. For each dataset, networks were trained on the training set for a total of 100 epochs with a batch size of 25 for GCNNs and 100 for 2D CNNs. After each epoch, the networks were tested on the validation set in order to select the best network configuration according to the lowest loss. After training, all networks were evaluated on the test set using their best configuration as described previously. The performance of each network was evaluated using average classification accuracy, precision, recall, F1 and area-under-ROC curve (AUC). Results associated with our protein classification experiments are shown in Table 3.

**Table 3:** Predictive performance of each network on the 4 classification tasks using average classification accuracy, precision, recall, F1 and area-under-ROC curve (AUC). Also shown is the computational performance measured as the number of training examples ingested by the networks per second. Best values per dataset per metric are highlighted in boldface.

| Model | Dataset | Examples/Sec | Loss | Acc | Prec | Recall | F1 | AUC |
| --- | --- | --- | --- | --- | --- | --- | --- | --- |
| 2DCNN | KRas/HRas | 531.75 | 0.0002 | 1.0 | 1.0 | 1.0 | 1.0 | 1.0 |
| GCNN | KRas/HRas | **835.00** | **1.08E-05** | 1.0 | 1.0 | 1.0 | 1.0 | 1.0 |
| 2DCNN | TSG/OG | **531.75** | 0.1740 | **0.9482** | **0.9538** | **0.9458** | **0.9498** | **0.9482** |
| GCNN | TSG/OG | 404.75 | **0.0326** | .9266 | 0.9259 | 0.9174 | 0.9216 | 0.9261 |
| 2DCNN | Kinase | **531.75** | **0.0683** | 0.9762 | 0.9654 | 0.9840 | 0.9746 | 0.9768 |
| GCNN | Kinase | 436.11 | 0.0710 | **0.9777** | **0.9655** | **0.9872** | **0.9762** | **0.9783** |
| 2DCNN | Enzyme | **531.75** | 1.2008 | **0.7804** | **0.8980** | **0.9129** | **0.9054** |  |
| GCNN | Enzyme | 320.68 | **1.1440** | 0.6018 | 0.8232 | 0.8022 | 0.8126 |  |

This table shows that the 2D CNN method proposed by Zacharaki et al. [47] and our GCNN method perform equally well for the K-Ras/H-Ras dataset on all metrics except for loss where GCNN performs better. GCNN performs better than the 2D CNN method for the Kinase dataset on all metrics except for loss. On the other hand, the 2D CNN method outperforms GCNN on all metrics except for loss on the oncogenes and enzymes datasets.

### 3.2 Computational Performance

Network models were implemented in Python3 using GPU-based Tensorflow 1.5 [65]. All networks were trained using a NVIDIA Quadro M4000 GPU with 8 GB of RAM. To measure the computational efficiency of our approach, we benchmarked the number of examples processed per second for each network variant. Table 3 includes a column describing this computational performance measured as the number of training examples ingested by the networks per second. This table shows that 2D CNN is computationally more efficient than GCNN on all datasets except for K-Ras/H-Ras, when using this metric.

## 4. Interpretability

A current issue in applying deep learning to scientific domains is the inability of interpreting what the models have learned during training. Such information may provide insightful features that help domain scientists understand the characteristics of the input data. Deep learning models are currently considered ‘black box’ algorithms and methods must be developed in order to identify what is happening ‘under the hood’. Several techniques such as gradient-based attribution [66], filter visualization [67], and layer activation maximization [68] have been used as methods of localizing learned parameters to the model’s input space. For example, attribution maps of 3D or 2D CNN models can be used to identify regions of the image that highly contribute to the final prediction of the model.

While this approach is somewhat useful in debugging network training to see if networks are learning desired features, it is not sufficient to tie salient regions in 3D or 2D images to specific biologically meaningful substructures. One of the benefits of GCNNs over CNNs is the ability to compute attribution in respect to residues instead of regions in space. This allows for a more interpretable representation formatted in a way that is also more meaningful to biologists who typically map primary onto tertiary structure to describe important characteristics of a protein. We applied attribution interpretation techniques on our GCNN models on the K-Ras/H-Ras dataset in order to 1) validate our network’s ability to abstract features from the data and 2) determine if the network is focusing on biologically meaningful substructures.

### 4.1 Ras Proteins

The Ras protein is frequently mutated in about 20% of all human cancers [51] and it is the subject of significant research due to the difficulty in designing efficient inhibitors. There are 3 Ras isoforms, K-Ras, H-Ras and N-Ras. Although there are 2 isoforms within the K-Ras class, K-Ras4A and K-Ras4B, which arise from alternative RNA splicing, we only consider the latter for this study and referred to it as K-Ras [50]. Ras proteins share 90% sequence identity, with conserved structural properties. However, isoform-specific residue differences, which occur in the second half of the sequence, are believed to play a role in their structural differences. Understanding the details of the specific characteristics of each class and how they can be exploited to design better drugs, is at the center of cancer research [50, 51]. In this study we focus on the K-Ras and H-Ras subtypes. Our results are preliminary and illustrate the potential of our method to target this important problem.

### 4.2 Attribution Method

#### 4.2.1 Gradient-Based Attribution

We use gradient-based attributions methods to determine how features along the protein structure are contributing to the classification. The attribution method we used is “gradient*input” and is defined as

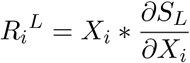

where *X* is the input features, *S* is the loss function, *L* is the class label. The attributions were calculated over both input tensors *V* and *C*, using soft-max cross-entropy as the loss function. After this, the sum of each feature contribution along the feature axis for both tensors was calculated, resulting in a single feature-level attribution tensor *R* ∈ ℝ^*Nx*1^. The attributions were then min-max normalized providing an interpretable scale of node importance with −1.0 indicating negative evidence towards the classification and 1.0 indicating positive evidence towards the classification. This method allows us to find significant contributions at the residue level. In the next section, we describe a method to identify substructures or segments of residues with similar attribution.

#### 4.2.2 Finding Attributed Substructures

To find statically significant substructures, we processed the attributions using Bellman k-segmentation [69] to cluster residue contribution along the primary structure of the protein. This algorithm generates a segmented constant-line fit to a data series, and helps in reducing noise present in the attributions in order to facilitate analysis. Given a desired number of segments *k*, the algorithm returns both residue membership for each segment and the average contribution of the segment.

For purposes of interpreting our model with respect to the K-Ras/H-Ras dataset, we selected a *k* value of 25. This value is inline with the number of unique structural partitions, which were referenced from Mattos et al.’s studies [50, 51]. These partitions include the common underlining secondary structures of Ras as well as relevant regions and evolutionary conserved substructures.

#### 4.2.3 Determining Class-Significant Features

In order to find which substructures are the most significant to the classification of K-Ras and H-Ras, we first found the average attribution between all unclustered examples for each class respectively. K-segmentation was then applied to the average class attribution to find the most significant substructures of each class. Since we are only interested in finding substructures which 1) are highly indicative of their parent class and 2) are most differentiating from the opposing class, we calculated the union of these two conditions using the following

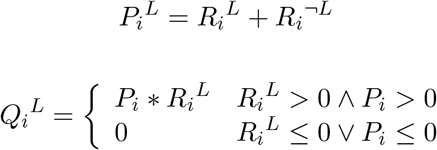

where *P*^*L*^ are the areas which provide the most differentiation and *Q*^*L*^ are those areas in *P*^*L*^ which positively contribute to the structures class membership. Here, we assume that areas with overall positively contribution between both classes are those in which the network had the most ease in finding differentiating features.

### 4.3 Interpreting Attributions

#### 4.3.1 Validating Generalization

We calculated the attribution along the sequence of amino acids for all structures in the K-Ras/H-Ras dataset. The attributions were first aligned by sequence and then sorted according to the lowest loss for both classes separately. Figure 2 shows the top 20 examples from unique PDBs for both classes ordered in descending order according to loss, with rows representing the proteins labeled according to their PDB ID and columns representing the residues in the sequence. Positively contributing residues are colored in red and negatively contributing residues are colored in blue. Notice that there are 2 rows per protein, one that shows the residues in red or blue color according to their contribution and another that shows the gaps or missing residues in the corresponding PDB file in dark red color.

**Figure 2:**
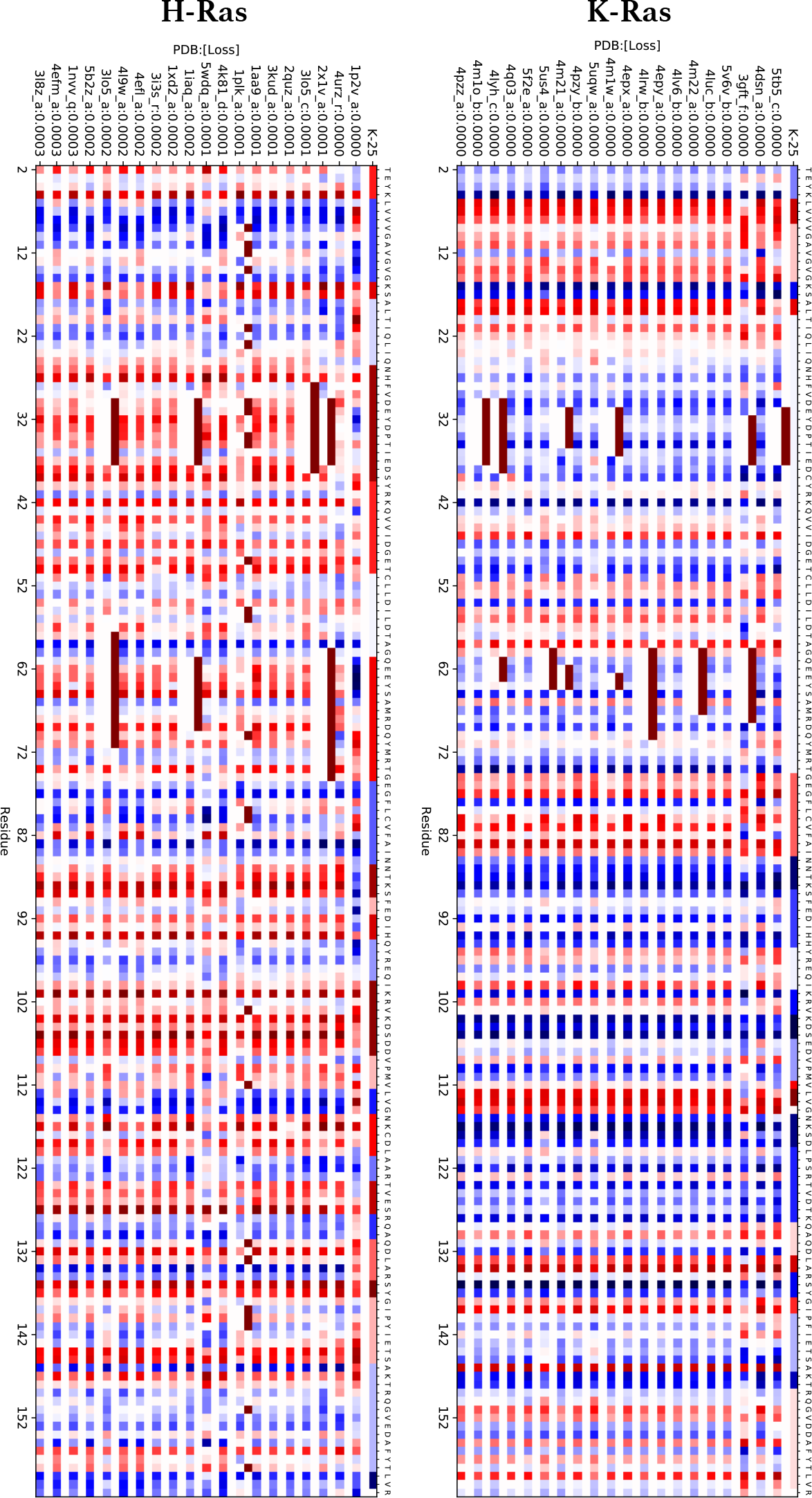
Plots of residue attributions for the top 20 lowest-loss examples from the K-Ras and H-Ras datasets. Each example corresponds to a protein which is labeled according to its PDB ID. Thus, the rows correspond to either H-Ras or K-Ras proteins and the columns correspond to the residues 2-160 in the sequence. They are aligned starting at residue T2. Each attribution is preceded with a row indicating missing residues in the PDB with dark red color. Residue contribution is colored on a blue to red scale with blue indicating negative contribution and red indicating positive contribution. To find statically significant substructures, we clustered the attributions using Bellman k-segmentation. The first row of each plot shows the 25 most significant clusters along all examples in each dataset respectively.

There are a few important things to note from this visualization. First, the commonality between attributions within a class is a good indicator that the GCNN method is learning abstracted features, and not simply memorizing each example. Second, we can also observe clear differences between attributions from the two classes. Lastly, while containing some noise between examples, the attributions appear to form clusters around specific areas along the sequence. These clusters can be more easily seen in the top row of each plot which shows the 25 most significant clusters between all examples in each class respectively. While providing a good validation that the network is generalizing and learning abstracted features, it is hard to pinpoint exactly how the clusters correspond to substructures in the proteins using this visualization without prior understanding of how structural features map onto primary structure. We address this issue in the next subsections.

#### 4.3.2 Visualizing Class Features

To better identify the features that are being referenced by the attributions, we also visualized the overall class attributions overlaid on 3D renderings of the proteins. Figure 3 shows 10 overlaid examples from each class rendered using PyMOL [70]. The coloring in this figure is used to highlight the 25 significant clusters discussed in the previous section and depicted in the top row (K-25) of the K-Ras and H-Ras attribution plots in Figure 2. This visual suggests that attribution values when rendered on 3D representations can be used to identify structural differences between classes. More importantly, this visual allows us to see how the segments derived from the attributions match many of the substructures of the Ras protein family and how they align with important sequence or structural differences between K-Ras and H-Ras.

**Figure 3:**
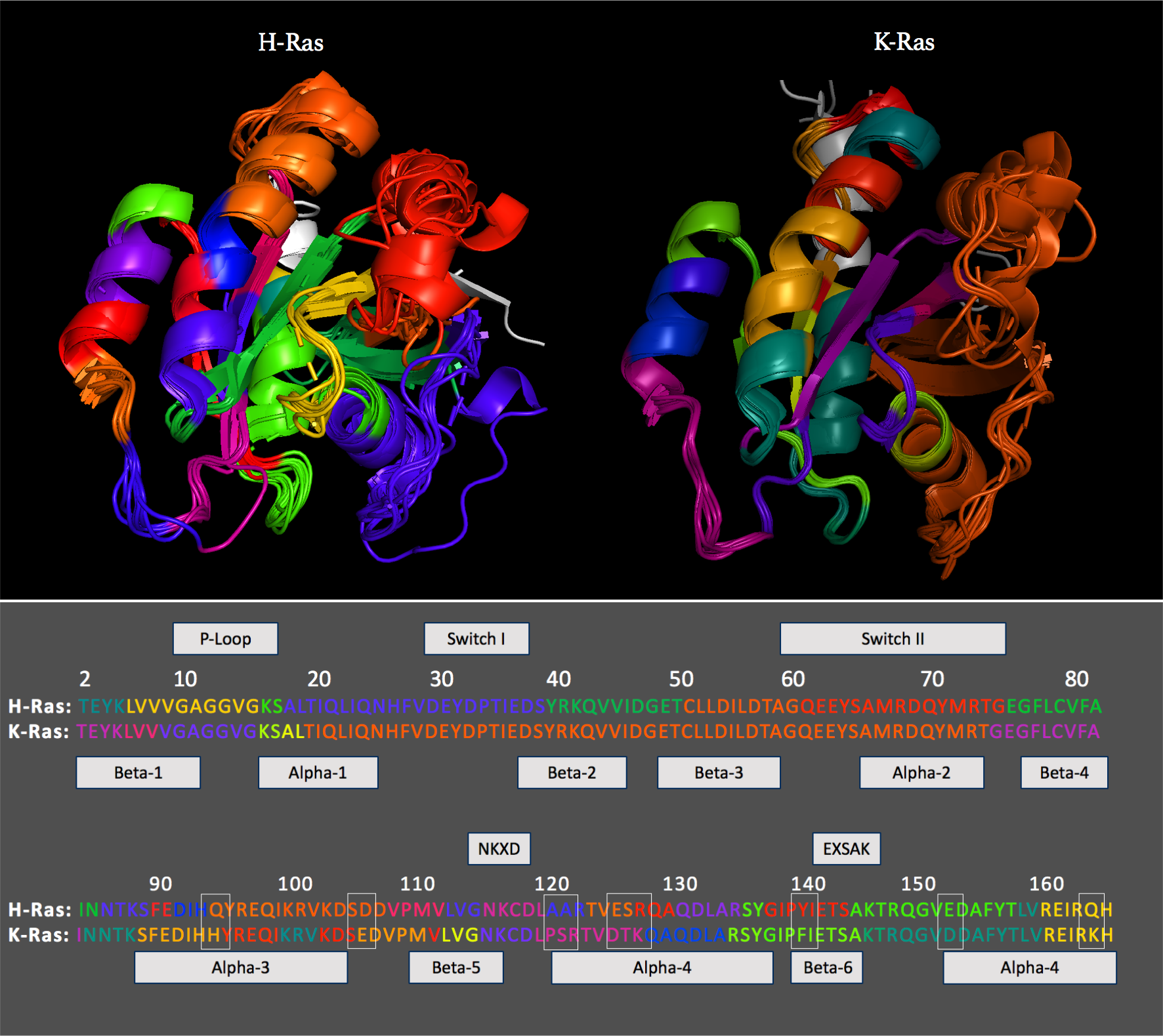
PyMol rendering of 10 overlayed examples from both H-Ras (left) and K-Ras (right). Structures are colored as to help in identifying the 25 distinct substructures derived from analysis of the network’s attribution, which are shown in the top row (K-25) of the attribution plots in Figure 2. The bottom portion of the figure displays the primary sequence of H-Ras and K-Ras colored to match the PyMOL renderings. Above them are labels for important Ras fragments aligned to the sequences and below them are labels for the Ras secondary structures. Residues that are different between K-Ras and H-Ras are boxed in white.

#### 4.3.3 Biological Significance

Figure 4 shows 10 overlaid structures corresponding to H-Ras (left) and K-Ras (right) isoforms. These structures are colored according to feature significance with respect to its class, with areas highlighted in darker red indicating more importance according to the network’s attribution. The figure identifies five important substructures within Ras which include the active sites’ switch I, switch II, and P-loop and the secondary structures alpha-helix 3 and 4. The red-colored areas of H-Ras correspond to switches I and II and the last portion of helix 3. There is an interplay between these regions that determines conformational changes in going from T state to R state as indicated in [50]. In the T state, helix 3 clashes with switch II, promoting a disordered conformation manifested in switch II, whereas in the R state there is room for the switch II helix to form. According to [50], K-Ras isoforms appear to favor the T state more so than H-Ras, which means that helix 3 tends to be more straight in H-Ras than in K-Ras. Notice that this helix tends to kink at residue 95 which is different in both isoforms and it is also highlighted in red in the H-Ras proteins depicted in Figure 4. Another highlighted area among the H-Ras proteins is the top portion of helix 4, which may be affected when helix 3/loop 7 shift toward helix 4 as part of the state R conformation.

There are fewer highlighted areas identified in the K-Ras proteins depicted in Figure 4. We believe that a possible explanation is that the dataset has a significantly larger number of H-Ras proteins in comparison with K-Ras. Residue 132 in helix 4 is among the positively attributed for the K-Ras proteins as well as the strand beta 6. We have not been able to tie these attributions to biological ones. However, there are other highlighted areas such as the last portion of beta 1 and the first portion of helix 1 which surround the P-loop. Also the last portion of beta 1 corresponds to residue 12 which is mutated in many Ras proteins and it is highly relevant to cancer research. Finally, the last portion of switch II is also highlighted in the K-Ras proteins.

**Figure 4:**
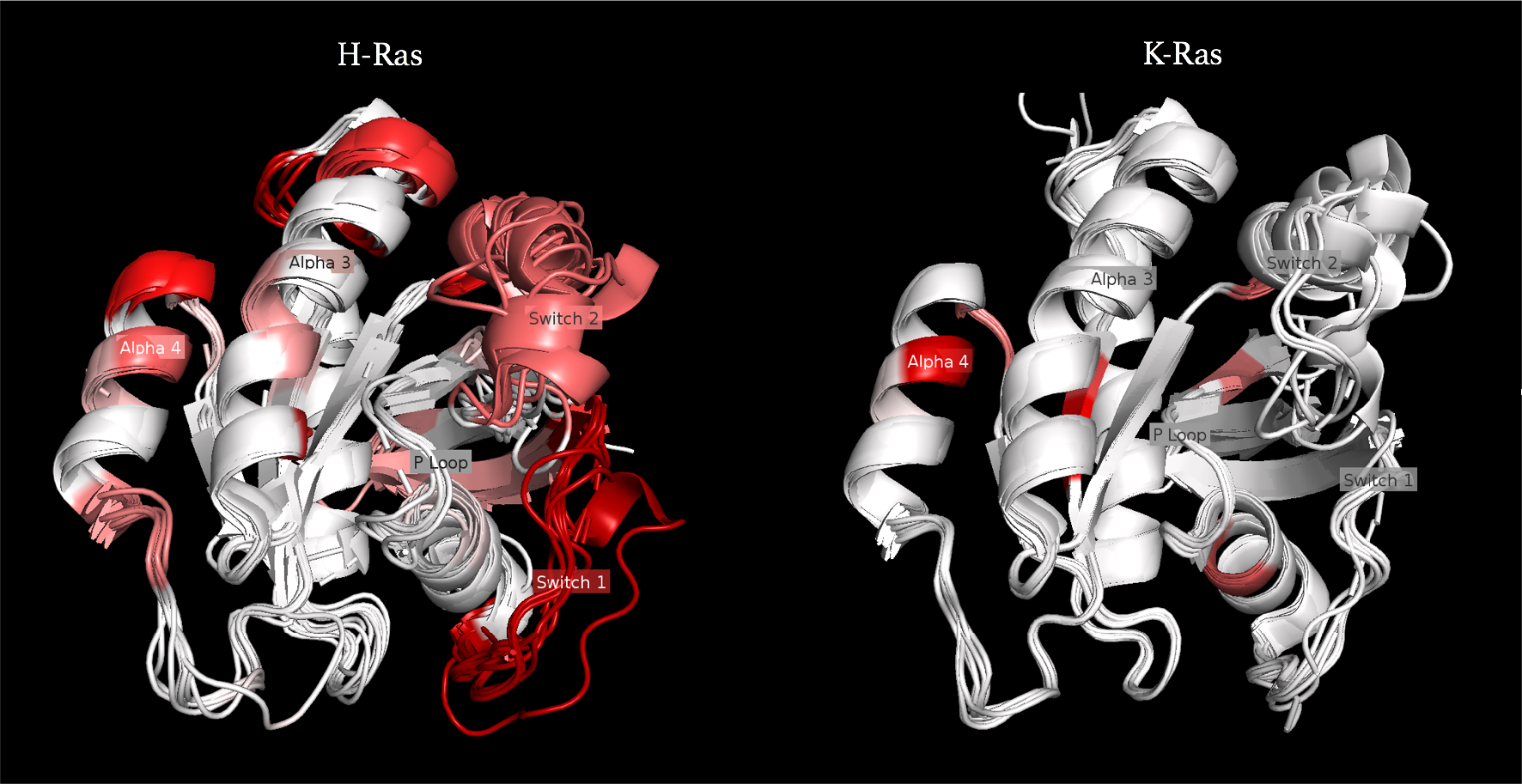
PyMOL rendering of 10 overlaid examples from both H-Ras (left) and K-Ras (right). Structures are colored according to feature significance with respect to its class, with areas highlighted in darker red indicating more importance according to the network’s attribution. Five important substructures to Ras are labeled including the active sites’ switch I, switch II, and P-loop and the secondary structures alpha helix 3 and 4.

## 5. Conclusion

### 5.1 Discussion

In this work, we discuss the application of CNNs to the analysis of molecular structures. We discuss the most commonly used approaches, 3D CNNs and 2D CNNs, and analyze their strengths and weaknesses. We present an alternative approach, GCNN, that provides a realistic spatial representation of molecular structures without the cost associated with using 3D volumes. The proposed methodology to construct protein graph representations encodes spatial relationships between residues. Also, we define a spatial graph convolution strategy that can best take advantage of the dense graph representation. We apply GCNNs to the classification of protein structures.

We compare the results produced by our GCNN to those produced by the 2D CNN approach presented by Zacharaki et al. [47], which to be best of our knowledge represent the best results from a deep learning method found in the literature. This comparisons show that our GCNN approach performs comparably to the 2D CNN approach in terms of predictive performance but it is computationally more expensive than the 2D CNN approach.

The graph-based approach has significant advantages over the 2D CNN or 3D CNN approaches. First, it can be transferred to other structural biology applications such as the protein-ligand binding affinity prediction, which is a regression problem. Second, as stated before, it is more efficient than the 3D CNN approach both in terms of computational cost and storage space. Third, unlike the 2D CNN and 3D CNN approaches, it provides great potential for interpretability. We explore the latter by focusing on the K-Ras/H-Ras dataset because understanding the structural differences between these isoforms is currently an active area of research and detailed descriptions of biologically meaningful differences can be found in the literature.

Our results show that the GCNN learns a set of features to make the classification that can be traced back to regions that biologists have identified in the Ras proteins. These networks separate the proteins in segments, some of which are in correspondence with secondary structures or biologically relevant loops and switches, or different residues, and then identify those segments that are relevant to the distinction between classes. These findings are remarkable and suggest that it is possible to learn from these black boxes when they are fed with interpretable data representations and analyzed with appropriate methods. We hope that this study starts a new trend in the application of deep learning to biological data that does not stop at the classification or regression results but digs deeper into information extraction. Although these technologies are undoubtedly powerful, there is still a great deal of work to be done to achieve their full potential for scientific discoveries.

### 5.2 Further Work

Although the proposed approach scales considerably better than the 3D CNN approach, there are still a number of computational issues that need to be addressed in order for this new methodology to be suitable for large-scale training. The main issue is scaling graph convolutions to graphs that contain more than 1,000 nodes. This issue can be addressed through a model parallel implementation of graph convolutions, which are able to partition the total graph over a number of available GPUs.

Predictive quality of GCNN models may also be improved by providing a biologically-aligned augmentation strategy using molecular dynamic simulation to generate valid training examples. Augmented examples can be generated prior to model training and managed with easy thanks to the GCNNs lean input data representation. This approach would require a large amount of computational resources to provide dataset sizes that are more inline with the current state of the art deep learning methods which are typically 3 to 4 degrees of magnitude greater than the datasets used in this study.

While this paper mainly focused on introducing an efficient and interpretable method for protein classification tasks, we plan to apply this method to the protein scoring and QA problems which are important components of the protein structure prediction pipeline [32]. The performance of DeepMind’s AlphaFold [71], during the recent CASP13 (the 13th Critical Assessment of techniques for protein Structure Prediction) competition [72] has shown the enormous potential of applying deep learning methodologies to the field. We are also working on applying this method the prediction of protein-ligand binding affinity, but more work is required to format our architecture on atomic-level data.

Also, we are considering a number of variations of gradient-based attribution commonly used for 2D images, specifically integrated-gradient attribution [66], to reduce noise. Integrated-gradient attribution has been shown to provide higher-fidelity attributions for CNNs, but work needs to be conducted to format this approach for our GCNN architecture. Finally, we plan to investigate the optimal design and assembly of a minimalistic network that allows us to achieve similar or better results at minimal cost. This would help in providing higher interpretability as the number of parameters in the network would be less. Our current results show that GCNNs can function at comparable levels to powerful ‘data-agnostic’ approaches like CNNs with reduced number of trainable parameters.

## Acknowledgments

This work was supported by the Director, Office of Science, Office of Basic Energy Sciences, of the U.S. Department of Energy under Contract No. DE-AC02-05CH11231. The authors would like to acknowledge Prof. Xinlian Liu and Thomas Corcoran for valuable discussions. Also the authors would like to acknowledge Prof. Nelson Max of University of California, Davis, and Drs. Peter Zwart, Steven Farrell, and Paolo Calafiura of Lawrence Berkeley National Laboratory for their insight and constructive feedback. Computing allocations were provided through the Cori supercomputer at the National Energy Scientific Computing Center (NERSC), the CAMERA GPU system at LBL, and the Exalearn initiative GPU system at LBL.

## Author Contributions

RZR created, designed, developed, and ran the models, created the images, and wrote the paper. SC conceived and directed the project and contributed to the writing of the paper.

